# In situ 10-cell RNA sequencing in tissue and tumor biopsy samples

**DOI:** 10.1101/444182

**Authors:** Shambhavi Singh, Lixin Wang, Dylan L. Schaff, Matthew D. Sutcliffe, Alex F. Koeppel, Jungeun Kim, Suna Onengut-Gumuscu, Kwon-Sik Park, Hui Zong, Kevin A. Janes

## Abstract

Single-cell transcriptomic methods classify new and existing cell types very effectively, but alternative approaches are needed to quantify the individual regulatory states of cells in their native tissue context. We combined the tissue preservation and single-cell resolution of laser capture with an improved preamplification procedure enabling RNA sequencing of 10 microdissected cells. This in situ 10-cell RNA sequencing (10cRNA-seq) can exploit fluorescent reporters of cell type in genetically engineered mice and is compatible with freshly cryoembedded clinical biopsies from patients. Through recombinant RNA spike-ins, we estimate dropout-free technical reliability as low as ~250 copies and a 50% detection sensitivity of ~45 copies per 10-cell reaction. By using small pools of microdissected cells, 10cRNA-seq improves per-cell reliability and sensitivity beyond existing approaches for single-cell RNA sequencing (scRNA-seq). Accordingly, in multiple tissue and tumor settings, we observe 1.5–2-fold increases in genes detected and overall alignment rates compared to scRNA-seq. Combined with existing approaches to deconvolve small pools of cells, 10cRNA-seq offers a reliable, unbiased, and sensitive way to measure cell-state heterogeneity in tissues and tumors.

## INTRODUCTION

Tumors are complex mixtures of cells that are heterogeneous in their genetics, lineage, and microenvironment (1,2). Whole-tumor profiles of genes and transcript abundances yield inter-tumor differences that are clinically important for patient prognosis (3-5), but these cellular profiles are population averages (6). The tumor microenvironment contains several different cell types that vary among cases (7-12). At the single-cell level, cancer cells are heterogeneous and genetic subclones evolve as the disease progresses (13,14). Tumor cells also display non-genetic heterogeneity and can switch between regulatory states in a reversible and context-dependent manner (15-17). Together, these variations dictate phenotypic differences such as proliferative index, metastatic potential, and response to therapy (16, 18-22).

Assessing intra-tumor heterogeneity of gene regulation requires precise transcriptomic measurements of a very small number of cells isolated from within the tumor context. The current methods for single-cell RNA sequencing (scRNA-seq) are powerful in their ability to profile thousands of individual cells and identify differences in genotype or lineage within a mixed population. However, the first step of most large-scale scRNA-seq methods is some form of tissue dissociation and single-cell isolation, which can alter transcriptional profiles and confound downstream analyses (23,24). Further, scRNA-seq methods struggle with technical variability, including "dropout" of medium-to-low abundance transcripts that yield zero aligned reads (25-28). The 3–20% conversion efficiency (25,26,29-31) of RNA to amplifiable cDNA is problematic given estimates that 90% of the transcriptome is expressed at 50 copies or fewer per cell (32). While valid for the most consistently expressed genes and markers within a sample, scRNA-seq data miss a large proportion of the transcriptome (32,33). Measuring single-cell expression profiles in situ is even more challenging because of losses incurred during biomolecule extraction as well as non-mRNA contaminants, which can be considerable in stroma-rich specimens. Collectively, these hurdles make it difficult to measure tumor-cell regulatory heterogeneities reliably and evaluate their functional consequences.

Multiple studies have reported a pronounced improvement in gene detection and technical reproducibility when using 10–30 cells of starting material rather than one cell (31, 34-39). The increased cellular RNA offsets losses incurred during reverse transcription, enabling more reliable downstream amplification. The gains are irrespective of amplification strategy and detection platform, and they are more dramatic than when increasing the starting material another tenfold to 100 cells. Previously, we combined the technical advantages of 10-cell pooling with the in-situ fidelity of laser-capture microdissection (LCM) to devise a random-sampling method called “stochastic profiling” (38,39). The method identifies single-cell regulatory heterogeneities by analyzing the statistical fluctuations of transcriptomes measured repeatedly as 10-cell pools microdissected from a cell lineage (38,40). Pooling improves gene detection and technical reproducibility; repeated sampling is used to extract single-cell information. Genes with bimodal regulatory states (41) create skewed deviations from a null model of biological and technical noise, which parameterize the underlying population-level distribution more accurately than single-cell measurements (36,42). By applying stochastic profiling to breast-epithelial spheroids and gene panels measured by quantitative PCR or microarray, we uncovered multiple regulatory states relevant to 3D organization and stress responses (18,43,44). However, this early work did not stringently evaluate the importance of sample integrity for primary tissues from animals or patients, nor did it involve probe-free measures of 10-cell data like RNA sequencing.

Here, we report improvements in sample handling, amplification, and detection that enable RNA sequencing of 10-cell pools isolated from tissue and tumor biopsies by LCM and its extensions. We find that cryoembedding of freshly isolated tissue pieces is crucial to preserve the localization of genetically encoded fluorophores in engineered mice used for fluorescence-guided LCM. By incorporating ERCC spike-ins at non-disruptive input amounts in the amplification, we calibrate sensitivity and provide a standard reference to compare with other scRNA-seq methods (45). Sample tagging and fragmentation (tagmentation) is accomplished by Tn5 transposase (46), which is compatible with the revised procedure as well as with past 10-cell amplifications. We sequence archival samples that had previously been measured by BeadChip microarray to provide a side-by-side comparison of transcriptomic platforms with limiting material (38,47). Applying 10-cell RNA sequencing (10cRNA-seq) to various mouse and human cell types isolated by LCM, we obtain substantially better exonic alignments and gene coverage relative to prevailing scRNA-seq methods. The realization of 10cRNA-seq by LCM creates new opportunities for stochastic profiling (42) and other unmixing approaches (36) to deconvolve single-cell regulatory states in situ.

## MATERIALS AND METHODS

### Cell and tissue sources

The MCF10A-5E breast epithelial cell samples were described previously (38). KP1 small-cell lung cancer cells (48) were grown as spheroids in RPMI Medium 1640 with 10% FBS, 1% penicillin-streptomycin, and 1% glutamine. KP1 spheroids were pelleted and mixed in Neg-50 (Richard-Allan Scientific) before cryoembedding. *Cspg4-CreER;Trp53^F/F^;Nf1^F/F^;Rosa26-LSL-tdT* mice (49) were housed in accordance with IACUC Protocol #3955. Animals were administered 200 mg/kg tamoxifen by oral gavage for five days, and brains were harvested at 12 days or 183 days after the last administration. A labeled glioma arising the olfactory bulb at 165 days after the last tamoxifen administration was also used. Breast cancer samples were collected as ultrasound-guided core needle biopsies during diagnostic visits in accordance with IRB Protocol #19272. Each core biopsy was divided into multiple pieces before cryoembedding. Unless otherwise indicated, all samples were freshly cryoembedded in a dry ice-isopentane bath and stored at –80°C wrapped in aluminum foil.

### Cryosectioning

Samples were equilibrated to –24°C in a cryostat before sectioning. 8 μm sections were cut and wicked onto Superfrost Plus slides. To preserve fluorescence localization of tdT and EGFP, slides were precooled on the cutting platform for 15–30 sec before wicking, and the section was carefully placed atop the cooled slide with forceps equilibrated at –24°C. Then, the slide was gently warmed from underneath by tapping with a finger until the section was minimally wicked onto the slide. All wicked slides were stored in the cryostat before transfer to –80°C storage on dry ice. Frost buildup was minimized by storing cryosections in five-slide mailers.

### Staining, dehydration, and laser-capture microdissection

For cryosections lacking fluorophores, slides were stained and dehydrated as described previously (38,39). Briefly, slides were fixed immediately in 75% ethanol for 30–60 sec, rehydrated quickly with water, stained with nuclear fast red (Vector Labs) containing 1 U/ml RNAsin-Plus (Promega) for 15 sec, and rinsed two more times with water before dehydrating with 70% ethanol for 30 sec, 95% ethanol for 30 sec, and 100% ethanol for 1 min and clearing with xylene for 2 min. tdT- and EGFP-labeled cryosections were not stained and instead began with the 70% ethanol dehydration step that also provided solvent fixation. After air drying, slides were microdissected immediately on an Arcturus XT LCM instrument (Applied Biosystems) using Capsure HS caps (Arcturus). The smallest spot size was used, and typical instrument settings of ~50 mW power and ~2 msec duration yielded ~25 μm spot diameters capturing 1–3 cells per laser shot.

### RNA extraction and first-strand synthesis

RNA extraction and first-strand synthesis were similar to earlier protocols (38,39) with some minor modifications. HS caps were eluted for 1 hr at 42°C with 4 μl of digestion buffer containing 1.25x First-strand buffer (Invitrogen), 100 μM dNTPs (Roche), 0.08 OD/ml oligo(dT)_24_ with or without 5’-biotin modification (IDT), and 250 μg/ml proteinase K (Sigma). Samples containing ERCC spike-ins included a four-million-fold dilution of ERCC spike-in mixture 1 (Ambion). Eluted samples were centrifuged into 0.5 ml PCR tubes at 560 rcf for 2 min, the digestion buffer was quenched with 1 μl of digestion stop buffer containing 2 U/μl SuperAse-in (Invitrogen) and 5 mM freshly prepared PMSF (Sigma). 4.5 μl of the quenched extract was transferred to a 0.2 ml PCR tube, and reverse transcription was performed with 0.5 μl of SuperScript III (Invitrogen) for 30 min at 50°C followed by heat inactivation at 70°C for 15 min. Samples were placed on ice and centrifuged for 2 min at 18,000 rcf on a benchtop microcentrifuge.

### Streptavidin bead cleanup of biotinylated first-strand products

For 5’-biotin-containing samples, streptavidin magnetic beads (Pierce) were prepared in a 0.2 ml PCR tube on a 96S Super Magnet Plate (Alpaqua). Beads (6 μl per sample) were magnetized, aspirated, and resuspended in binding buffer (5 μl per sample) containing 1x First-strand buffer (Invitrogen), 4 M NaCl, and 0.02% (vol/vol) Tween-20. 5 μl of resuspended beads were added after first-strand synthesis, and samples were incubated for 60 min at room temperature with mixing every 15 min. Beads were pelleted on the magnet plate, resuspended in 100 μl high-salt wash buffer (50 mM Tris [pH 8.3], 2 M NaCl, 75 mM KCl, 3 mM MgCl_2_, 0.01% Tween-20). Beads were pelleted again on the magnet plate, and the pellet was washed once with 100 μl high-salt wash buffer. Next, beads were resuspended in 100 μl low-salt wash buffer (50 mM Tris [pH 8.3], 75 mM KCl, 3 mM MgCl_2_) and transferred to a fresh 0.2 ml PCR tube. Beads were pelleted again on the magnet plate, and the pellet was washed once with 100 μl low-salt wash buffer. After the last wash, the beads were resuspended in 5 μl 1x First-strand buffer for RNAse H treatment and poly(A) tailing.

### RNAse H treatment and poly(A) tailing

RNAse H digestion and poly(A) tailing were performed exactly as described previously (38,39). Briefly, template mRNA strands were hydrolyzed for 15 min at 37°C with 1 μl of RNAse H solution containing 2.5 U/ml RNAse H (USB Corporation) and 12.5 mM MgCl_2_. After RNAse H treatment, cDNA templates were poly(A)-tailed with 3.5 μl of 2.6x tailing solution containing 80 U terminal transferase (Roche), 2.6x terminal transferase buffer (Invitrogen) and 1.9 mM dATP. The tailing reaction was incubated for 15 min at 37°C and then heat-inactivated at 65°C for 10 min. Samples were placed on ice and spun for 2 min at 18,000 rcf on a benchtop centrifuge.

### Poly(A) PCR

Poly(A) PCR was carried out with several modifications to the earlier procedure (38,39). To each tailed sample, 90 μl of poly(A) PCR buffer was added to a final concentration of 1x ThermoPol buffer (New England Biolabs), 2.5 mM MgSO_4_, 1 mM dNTPs (Roche), 100 μg/ml BSA (Roche), 3.75 U Taq polymerase (NEB) and 1.5 U Phusion (NEB) and 2.5 μg AL1 primer (ATTGGATCCAGGCCGCTCTGGACAAAATATGAATTCTTTTTTTTTTTTTTTTTTTTTTTT). Each reaction was split into three thin-walled 0.2 ml PCR tubes and amplified according to the following thermal cycling scheme: four cycles of 1 min at 94°C (denaturation), 2 min at 32°C (annealing) and 2 min plus 10 sec per cycle at 72°C (extension); 21 cycles of 1 min at 94°C (denaturation), 2 min at 42°C (annealing) and 2 min 40 sec plus 10 sec per cycle at 72°C (extension). The tubes were cooled, placed on ice, and the reactions from three tubes for each sample were pooled and amplified according to the following thermal cycling scheme: five cycles of 1 min at 94°C (denaturation), 2 min at 42°C (annealing) and 6 min at 72°C (extension). Amplified samples were stored at −20°C until further use.

### Poly(A) PCR re-amplification

For sequencing, poly(A) PCR cDNA samples were reamplified as before (38,39) in a 100 μl PCR reaction containing 1x High-Fidelity buffer (Roche), 3.5 mM MgCl_2_, 200 μM dNTPs (Roche), 100 μg/ml BSA (Roche), 5 μg AL1 primer, and 1 μl of poly(A) PCR sample. Each reaction was amplified according to the following thermal cycling scheme: 1 min at 94°C (denaturation), 2 min at 42°C (annealing) and 3 min at 72°C (extension). The appropriate number of PCR cycles was determined by a pilot reamplification containing 20 μl of the PCR reaction above plus 0.25x SYBR Green monitored on a CFX96 real-time PCR instrument (Bio-Rad). The number of amplification cycles for each sample was selected to ensure that the reamplification remained in the exponential phase and there was sufficient cDNA for SPRI bead purification (typically 5–12 cycles).

### SPRI bead purification

Re-amplified samples were purified twice with 70% (vol/vol) Ampure Agencourt XP SPRI beads. SPRI beads were equilibrated to room temperature for 30 min, and 70 μl beads were added to the 100 μl reamplification product. After a 15-min incubation at room temperature, samples were magnetized for 5 min. The supernatant was removed with a gel-loading pipette tip, leaving ~5 μl volume in the well. Beads were gently washed twice on the magnet with 200 μl freshly prepared 80% (vol/vol) ethanol and aspirated with a gel-loading pipette tip. Residual ethanol was removed after the second wash, and beads were air-dried at room temperature for 10 min before resuspension in 10 μl elution buffer (10 mM Tris-HCl [pH 8.5]). Samples were magnetized at room temperature for 1 min, and the eluted supernatant was transferred to a new 0.2 ml PCR tube. The 10 μl elution was purified a second time with 7 μl beads and the same incubation, ethanol wash, and elution conditions as the first purification.

### RNA sequencing and analysis

Bead-purified cDNA libraries were quantified with the Qubit dsDNA BR Assay Kit (Thermo Fisher) using a seven-point standard curve and a CFX96 real-time PCR instrument (Bio-Rad) for detection. Samples were diluted to 0.2 ng/μl before tagmentation with the Nextera XT DNA Library Preparation Kit (Illumina) according to the manufacturer’s earlier recommendation to purify libraries with 180% (vol/vol) SPRI beads (Fig. S7). For each run, samples were multiplexed at an equimolar ratio, and 1.3 pM of the multiplexed pool was sequenced on a NextSeq 500 instrument with NextSeq 500/550 Mid/High Output v1/v2 kits (Illumina) at an average read depth of 4.2 million reads per sample (Fig. S6) or 7.5 million reads per sample (all others). Adapters were trimmed using fastq-mcf in the EAutils package (version ea-utils.1.1.2-537) with the following options: -q 10 -t 0.01 -k 0 (quality threshold 10, 0.01% occurrence frequency, no nucleotide skew causing cycle removal). Quality checks were performed with FastQC (version 0.11.7) and multiqc (version 1.5). Datasets were aligned to either the human (GRCh38.84) or the mouse (GRCm38.82) transcriptome along with reference sequences for ERCC spike-ins using RSEM (version 1.3.0) with the following options: --bowtie2 --single-cell-prior -- paired-end (Bowtie2 transcriptome aligner, single-cell prior to account for dropouts, paired end reads). RSEM read counts were converted to transcripts per million (TPM) by dividing each value by the total read count for each sample and multiplying by 10^6^. Mitochondrial genes and ERCC spike-ins were not counted towards the total read count during TPM normalization. The number of genes with TPM > 1 for each sample was calculated relative to the number of unique Ensembl IDs for the organism excluding ERCC spike-ins.

### Analysis of public scRNA-seq datasets

FASTQ files were downloaded from GSE75330, GSE60361, GSE103354 (plate-based), GSE66357, GSE113197, and PRJNA396019. FASTQ files were not available for the droplet-based dataset of GSE103354; therefore, BAM files were downloaded from SRR7621182 and converted to FASTQ format. Adapters were trimmed using fastq-mcf with the following options: -q 10 -t 0.01 -k 0 (quality threshold 10, 0.01% occurrence frequency, no nucleotide skew causing cycle removal). To compare with the other datasets, seqtk (version 1.3) was used to clip 15 bp unique molecular identifiers from the beginning of sequences in GSE60361 and GSE75330. All RNA-seq datasets were aligned to either the human (GRCh38.84) or the mouse (GRCm38.82) transcriptome as well as reference sequences for ERCC spike-ins, using RSEM with the following options: --bowtie2 --single-cell-prior (Bowtie2 transcriptome aligner, single-cell prior to account for dropouts). GSE113197 and PRJNA396019 also used --paired-end (paired end reads). TPM conversion and gene detection quantification were calculated as above.

### qPCR

For detection of specific targets in poly(A) PCR samples, qPCR was performed on a CFX96 real-time PCR instrument (Bio-Rad) as previously described (50). 0.1 μl or 0.01 μl of each preamplification was used with the qPCR primers listed in Supplementary Table S1. For relative quantification between ERCC spike-ins, qPCR amplicons were purified by gel electrophoresis, extracted, ethanol precipitated, and quantified by spectrophotometry. Purified amplicons were used to create a six-log standard curve based on ERCC amplicon copy number. All spike-ins were normalized to ERCC130 copy numbers to obtain relative abundance.

### Paired analysis of BeadChip microarrays and 10cRNA-seq

Microarray data (GSE120030) (38) were batch processed with the lumi R package (51) using a detection threshold of 0.05 and simple scaling normalization to obtain log_2_-normalized values that were converted to log_10_-normalized values. Gene names from the BeadChip files were merged to the extent possible with Ensembl IDs from the RSEM alignments by using HUGO Gene Nomenclature synonym tables to match current and retired gene names.

## RESULTS

Methods for profiling small quantities of cellular RNA have evolved considerably over the past decade, but they all involve the same fundamental steps: 1) cell isolation, 2) RNA extraction, 3) reverse transcription, 4) preamplification, and 5) detection (52). The original protocol for in situ 10-cell profiling combines LCM for cell isolation followed by proteinase K digestion for RNA extraction (39). The extracted material undergoes an abbreviated high-temperature reverse transcription with oligo(dT)_24_, and cDNA is carefully preamplified by poly(A) PCR (53) that generates sufficient 3’ ends for microarray labeling and hybridization (39) (Figure 1).

Unsurprisingly, the earliest steps in the procedure are the most critical for achieving the maximum amount of amplifiable starting material. To avoid losses, steps 1–4 (cell isolation through preamplification) are normally performed without intermediate purification. Therefore, buffers and reagents must be carefully tested and titrated to be mutually compatible throughout the “one-pot” protocol. Since description of the procedure (38,39), multiple commercial providers merged or were acquired, leading to the discontinuation of multiple RNAse inhibitors, the Taq polymerase, and the BeadChip microarrays. The collective disruptions in sourcing prompted a modernization of 10-cell profiling toward RNA-seq of primary material at a biopsy scale, including how tissue–tumor samples were handled before the start of the procedure (Figure 1).

**Figure 1.**
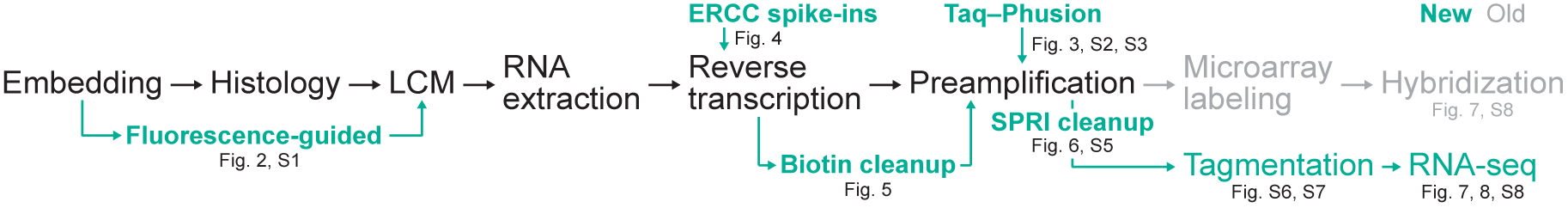
A revised transcriptomic pipeline for in situ 10-cell RNA sequencing. Substantive changes are indicated in green and gray.

### Protein localization for LCM requires fresh cryoembedding

To minimize extra handling steps that could degrade RNA, in situ profiling of clinical samples is ordinarily performed with rapid histological stains (38,52,54,55) (Figure 1). LCM can also be guided by fluorescence in place of histology when using cells or animals engineered to encode genetic labels (56,57). However, new challenges arise when seeking to preserve localization and brightness of encoded fluorophores during single-cell isolation and RNA extraction. Compared to polysome-bound mRNAs, fluorescent proteins diffuse much more readily, and chromophores may be damaged by the fixation and dehydration steps needed to preserve RNA integrity. Fluorescent-protein structure is preserved by chemical fixatives, but covalent crosslinking of biomolecules is unsuitable for extracting RNA from tissue. Fluorescence-guided profiling therefore entails a competing set of tradeoffs that must be balanced for optimal performance.

We reasoned that the greatest flexibility would be afforded by reporter mice expressing tandem-dimer Tomato (tdT)—a bright, high molecular-weight derivative of DsRed (58). Key handling parameters were evaluated using *Cspg4-CreER;Trp53^F/F^;Nf1^F/F^;Rosa26-LSL-tdT* mice, a model of malignant glioma (49). In these animals, administration of tamoxifen elicits sparse labeling of oligodendrocyte precursor cells (OPCs) in the brain, enabling fluorescence retention to be assessed in single cells. Extensive optimization of cryosectioning and wicking conditions was required to preclude fluorophore diffusion while ensuring reliable LCM pickup (see Methods). We found that an accelerated 70-95-100% ethanol series (38,39) maintained tdT fluorescence and localization of labeled cells through xylene clearing and dehydration (Figure 2A). Separately, using freshly embedded tissue from a “mosaic analysis of double markers” (MADM) animal that labels various brain lineages with EGFP, tdT, or both (59,60), we confirmed that EGFP fluorescence was also acceptably retained with the 70-95-100% ethanol series (Supplementary Figure S1). Although EGFP diffusion was noticeably greater compared to tdT owing to its smaller size (~28 kDa vs. ~54 kDa), we could nonetheless reliably identify the cell bodies of single EGFP-positive cells for LCM. Surprisingly, we found that fresh-tissue embedding was critically important for preserving single-cell localization and brightness. Snap-freezing before cryoembedding caused considerable loss and delocalization of tdT fluorescence, even when prefrozen material was rapidly embedded in dry ice-isopentane (–40°C) (Figure 2B,C). For mechanically challenging tissues in which embedding support is important for cryosectioning, we conclude that fresh-tissue embedding is essential for maximum biomolecular retention and integrity.

**Figure 2.**
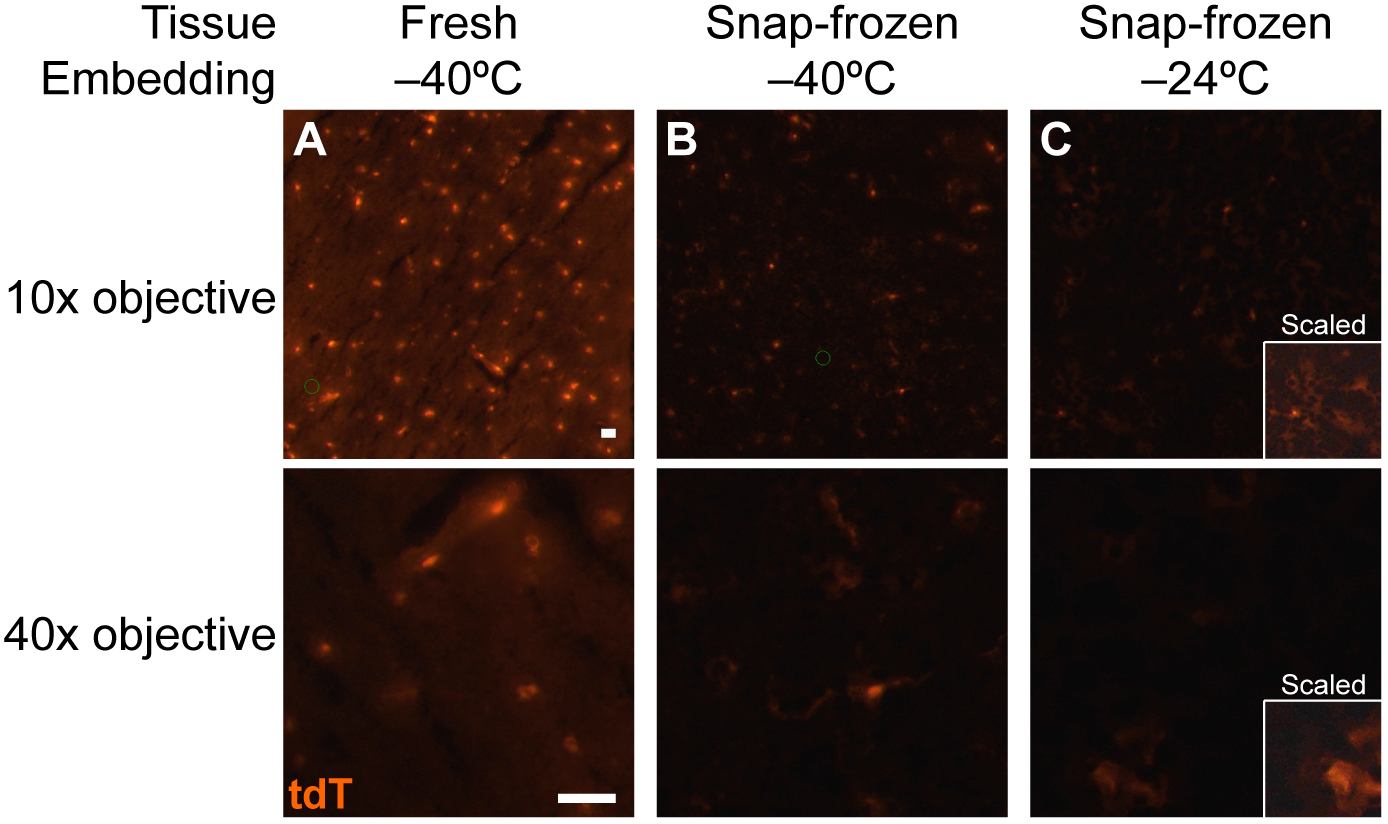
Fresh cryoembedding preserves tandem-dimer Tomato (tdT) fluorescence and localization better than snap-frozen alternatives. Brain samples from *Cspg4-CreER;Trp53^F/F^;Nf1^F/F^;Rosa26-LSL-tdT* animals were **(A)** freshly cryoembedded in Neg-50 medium with dry ice-isopentane (–40°C), **(B)** snap-frozen in dry ice-isopentane and then cryoembedded, or **(C)** snap-frozen and slowly cryoembedded in a cryostat (–24°C). Low- and high-magnification images were captured with the factory-installed color camera on the Arcturus XT LCM instrument. Images were exposure matched and are displayed with a gamma compression of 0.67. Insets have been rescaled to emphasize tdT diffusion away from the cell body. Scale bar is 25 μm.

### Improving poly(A) preamplification for modern RNA-seq

Previously, in situ 10-cell profiling was optimized for quantification by BeadChip microarray (38,39), but microarrays have been supplanted by RNA-seq for unbiased measures of the transcriptome (61) (Figure 1). An advantage of RNA-seq is that nucleic acids are detected regardless of origin, enabling use of exogenous RNA standards to calibrate sensitivity and quantitative accuracy when spiked into a biological sample (62-64). The versatility of RNA-seq is also a caveat, because all nucleic acids in a sample will be sequenced, including unwanted preamplification byproducts and contaminating DNA from mitochondria or the nucleus (65-67). In the original scRNA-seq report that used a variant of poly(A) PCR, only 37 ± 9% of sequenced reads aligned to RefSeq transcripts (68), and exonic alignment rates below 50% remain common (69). Therefore, we focused improvements to poly(A) preamplification towards ensuring that most sequencing reads aligned to the 3’ ends of cellular mRNAs.

In poly(A) PCR, cDNA is 3’ adenylated and then preamplified with a universal T24-containing primer called AL1 (53). We previously found that the amount of AL1 strongly influenced overall sensitivity of gene detection, with improvements noted at concentrations as high as 25 μM (39). Excess AL1 also drives nonspecific amplification of low molecular-weight primer concatemers (70), which do not influence gene measurements by quantitative PCR or microarray but create overwhelming contamination for RNA-seq. To improve poly(A) PCR, we screened a range of commercial Taq and proofreading polymerases along with empirical blends of those that maximized the intended ~500 bp cDNA products relative to nonspecific concatemer. We obtained a better-than-additive preamplification by combining Taq and Phusion polymerases (see Methods). An equal mixture of the two enzymes dramatically increased the yield of ~500 bp preamplification products relative to nonspecific concatemer (Figure 3A, lower). The empirical blend also significantly improved the preamplification of both high-abundance (*GAPDH*) and low-abundance (*PARN*) targets as measured by quantitative PCR (Figure 3A, upper). The two-enzyme blend further enabled a 10-fold decrease in AL1 primer concentration without detectable loss in preamplification efficiency (Figure 3B). The Taq-Phusion combination was superior for a primary breast-cancer biopsy (Figure 3) as well as two murine tissue sources: a murine small-cell lung cancer line derived from *Trp53^∆/∆^Rb^∆/∆^* lung epithelium (48) and tdT-labeled OPCs (Supplementary Figure S2 and S3), illustrating its generality. The enzyme modification created a viable starting point for combining poly(A) PCR preamplification with RNA-seq.

**Figure 3.**
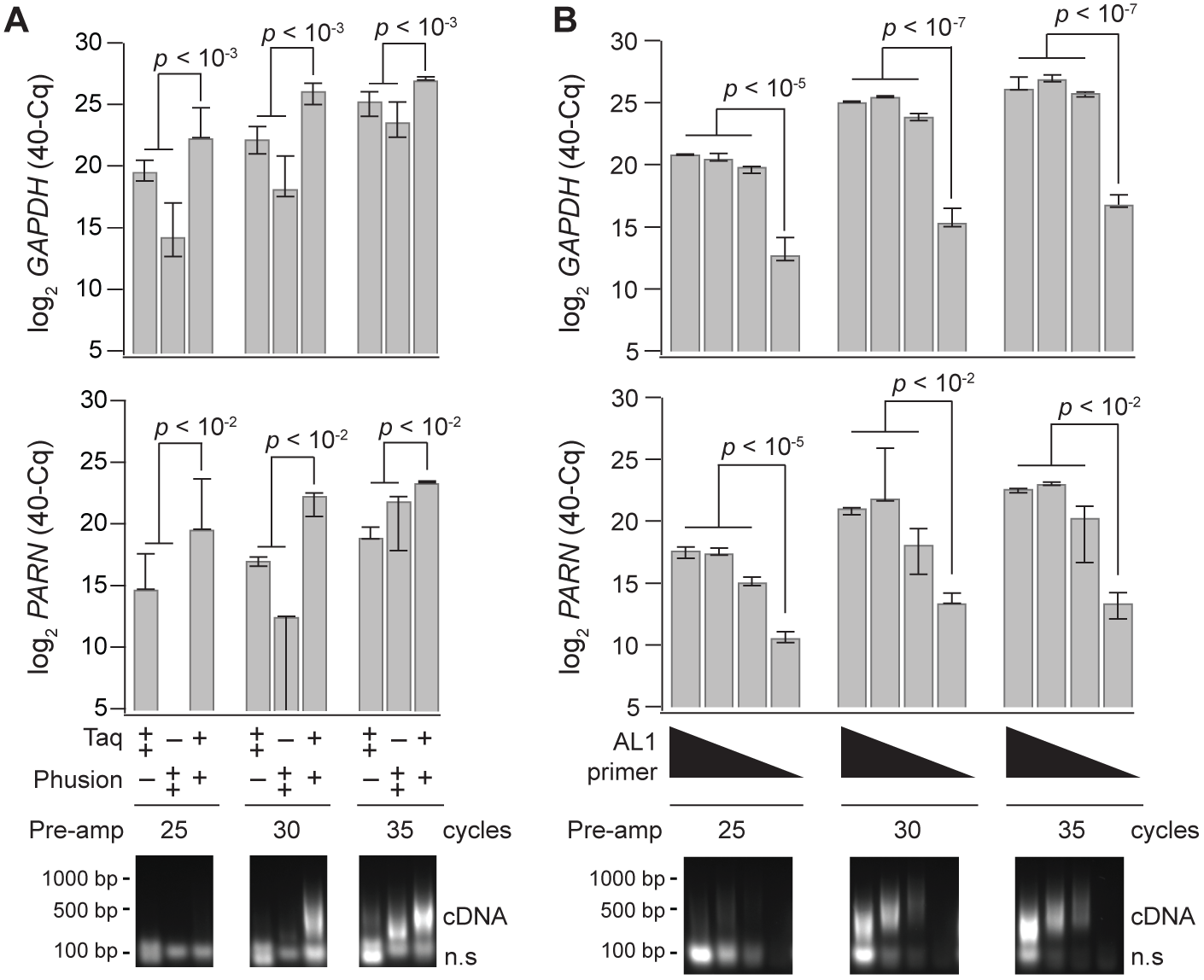
A blend of Taq–Phusion polymerases improves selective poly(A) amplification of cDNA and reduces AL1 primer requirements. Cells were obtained by LCM from a human breast biopsy and split into 10-cell equivalent amplification replicates. **(A)** Poly(A) PCR was performed with 15 μg of AL1 primer with Taq alone (10 units), Phusion alone (4 units) or Taq/Phusion combination (3.75 units/1.5 units). **(B)** Poly(A) PCR was performed with either 25, 5, 2.5 or 0.5 μg of AL1 primer and the Taq–Phusion blend from (A). Above—Relative abundance for the indicated genes and preamplification conditions was measured by quantitative PCR (qPCR). Data are shown as the median inverse quantification cycle (40–Cq) ± range from *n* = 3 amplification replicates and were analyzed by two-way (A) or one-way (B) ANOVA with replication. Below—Preamplifications were analyzed by agarose gel electrophoresis to separate poly(A)-amplified cDNA from nonspecific, low molecular-weight concatemer (n.s.). Qualitatively similar results were obtained separately three times.

Sensitivity, accuracy, and precision of the updated poly(A) PCR approach were assessed using recombinant RNA spike-ins as internal positive controls (64). A dilution of ERCC spike-ins was defined that did not detectably perturb the measured abundance of endogenous transcripts in RNA equivalents from 10 microdissected cells (Figure 4A). After poly(A) PCR of the spike-in dilution plus 100 pg RNA (~10 cells), we measured the relative abundance of individual spike-ins, using quantitative PCR (qPCR) to eliminate RNA-seq read depth as a complicating factor. Purified qPCR end products served as an absolute reference of each spike-in for cross-comparison (see Methods). We observed good linearity across 22 spike-ins spanning an abundance of ~10^4^ (Figure 4B). Deviations, technical noise, and dropouts all increased considerably for spike-ins below ~250 copies per reaction, consistent with previous reports (25). This collective measurement uncertainty restricts interpretation of single-cell data to highly expressed transcripts, but 10-cell pooling reduces the threshold to ~25 copies on average per cell. With poly(A) PCR, we did not observe qualitative dropout in more than 50% of technical replicates for spike-ins as dilute as four copies per reaction (ERCC85; Figure 4B), indicating good sensitivity. RNA spike-ins do not mimic the characteristics of endogenous transcripts extracted from cells, but they can provide a common reference to benchmark preamplification methods for RNA-seq (45). These experiments indicated that the improved poly(A) preamplification was sufficiently reliable for unbiased profiling of 10-cell transcriptomes.

**Figure 4.**
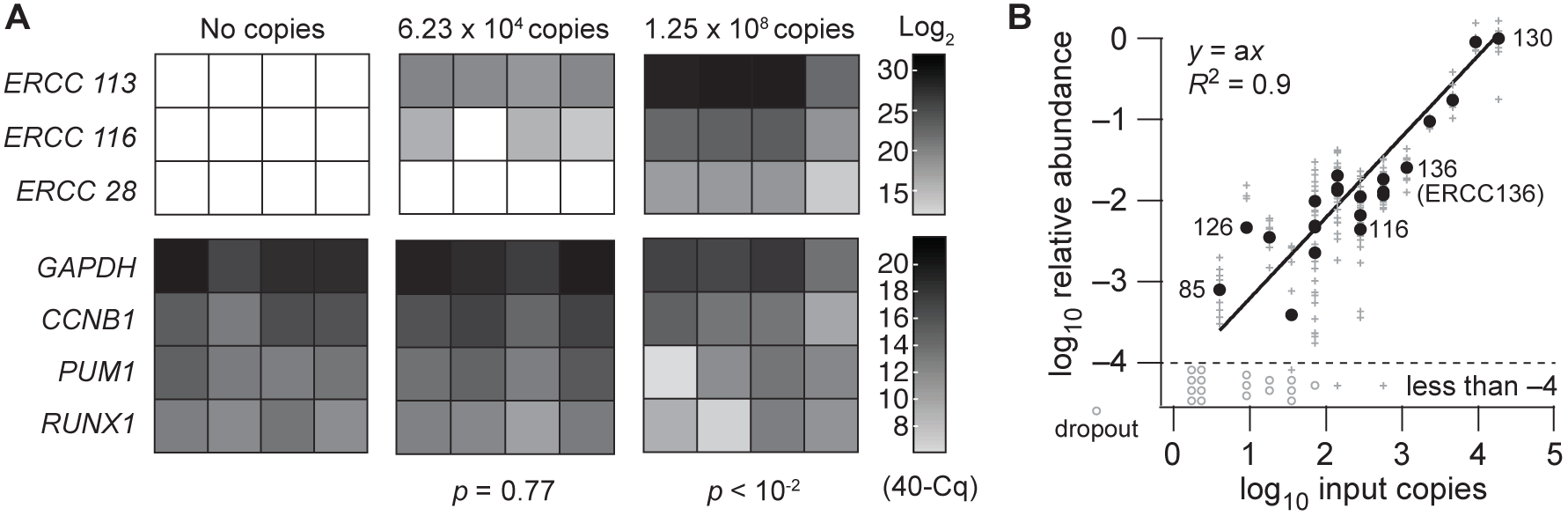
Optimized ERCC spike-in dilutions assess poly(A) PCR sensitivity and dynamic range without suppressing cDNA amplification of endogenous transcripts. **(A)** 100 pg RNA was supplemented with ERCC Mix 1 at the indicated dilutions and amplified via optimized poly(A) PCR. ERCC and endogenous gene abundances were measured by qPCR, and data are shown in grayscale as the inverse quantification cycle (40–Cq) from *n* = 4 amplification replicates. Negative effects of the ERCC spike-ins on endogenous genes (lower) were assess by two-way ANOVA with replication. **(B)** ERCC Mix 1 (6.23 × 10^4^ copies) was spiked into 100 pg RNA and amplified via optimized poly (A) PCR. Proportional abundance of ERCC standards was estimated with a seven-log dilution series from purified qPCR end products. Data are shown as the median 40–Cq (black) for 22 ERCC spike-in standards from *n* = 8 amplification replicates (gray) with undetected “dropouts” reported below (circles).

For RNA extraction from the LCM cap, an optimized digestion buffer is used containing proteinase K to release mRNAs from precipitated ribosomes (38). Proteinase K also digests nucleosomes, which may cause elution of contaminating genomic DNA. In past and current analyses of human LCM samples preamplified ± reverse transcription, we never found genomic copies of genes amplified within ~0.4% of measured mRNA transcripts (∆Cq ≥ 8 for 16 genes measured in four human cell types, Supplementary Figure S4). For mouse tissues, however, genomic copies were more prevalent and variable, with some genes measured as abundantly without reverse transcription as with it (Figure 5A and Supplementary Figure S4). Gel electrophoresis showed weak-but-detectable bands above the desired ~500 bp product in preamplifications without reverse transcription, implying nonspecific amplification (Figure 5A, lower). Concerned that the murine genome could compete with the amplification of cDNA, we appended an intermediate purification following reverse transcription with 5’-biotin-modified oligo(dT)_24_. Biotinylated cDNA was purified on streptavidin-conjugated magnetic beads, which could be separated from contaminants in the LCM extract and used as a starting template for poly(A) preamplification. Addition of the biotin cleanup step mildly improved the amplification of cDNAs and, importantly, eliminated the confounding abundance of murine genomic DNA (Figure 5B). We recommend biotinylated oligo(dT)_24_ and bead purification for mouse samples considering the recurrent challenges with genomic DNA (Supplementary Figure S4 and see Discussion).

**Figure 5.**
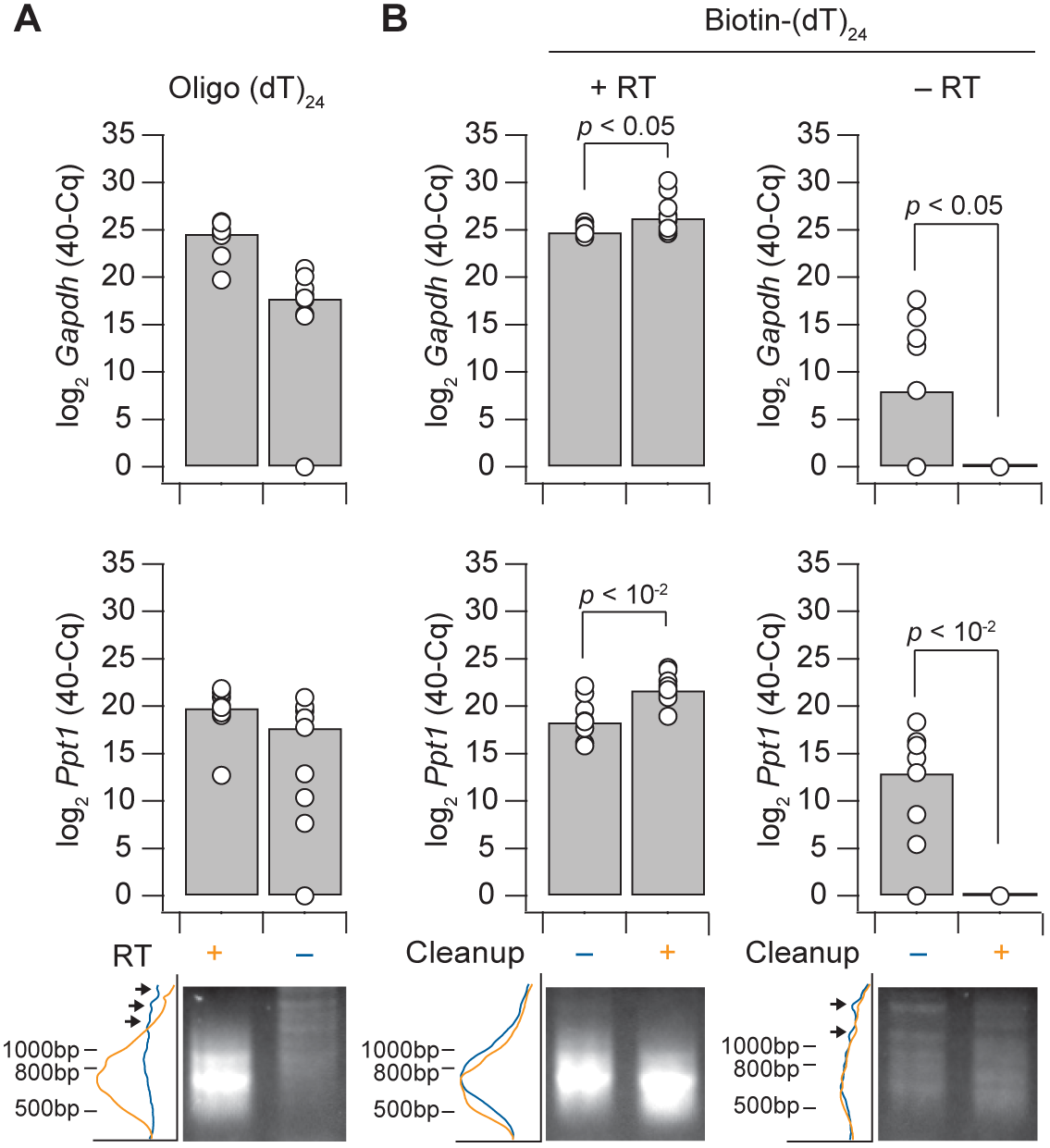
Poly(A) amplification of murine sequences without reverse transcription is eliminated with 5’-biotin-modified oligo(dT)_24_ and streptavidin bead cleanup. **(A)** Reverse transcription-free preamplification of genomic DNA confounds accurate quantification of some mRNAs. **(B)** Bead cleanup eliminates nonspecific preamplification of genomic DNA. Above—Data are shown as the median inverse quantification cycle (40–Cq, gray) of *n* = 3 independent experiments (three amplification replicates per experiment). Differences with and without bead cleanup were assessed by Wilcoxon rank sum test in MATLAB. Below—Preamplifications were analyzed by agarose gel electrophoresis to separate poly(A)-amplified cDNA from nonspecific, low molecular-weight concatemer (n.s.) and genomic amplification. Electrophoretic traces were analyzed by densitometry to the left of the image, with genomic amplicons highlighted (arrows).

Poly(A) PCR samples are kept dilute to avoid saturating the preamplification, but aliquots can be carefully reamplified up to microgram scale for hybridization (38,39). In preparing libraries for sequencing, we pursued tagmentation using Tn5 transposase because addition of sequencing adapters is sterically impossible within the ~40 bp distal ends of a PCR amplicon (71). The steric restrictions of Tn5 were advantageous for pruning away the long, A-repetitive universal primer from poly(A) amplicons that would otherwise be wastefully sequenced. Commercial Tn5 tagmentation kits (Nextera XT) require 1000-fold less material than past microarray hybridizations, prompting reevaluation of how the 10-cell libraries were prepared. We retained the mid-logarithmic reamplification approach described previously (38) but substituted paramagnetic Solid Phase Reversible Immobilization (SPRI) beads for library purification (72). Two rounds of purification with 70% (vol/vol) SPRI beads eliminated ~99% of primer dimers and concatemers in 10-cell reamplifications from various sources (Figure 6 and Supplementary Figure S5). Reamplified samples yielding at least 200 ng of purified product (Supplementary Figure S6) were tagmented at 1-ng scale according to the Nextera XT protocol. Although poly(A) amplicon sizes are centered at ~500 bp (Figure 1), we found that the higher SPRI bead ratio recommended for 300–500 bp inputs (180% [vol/vol] beads) was essential for purification of tagmented libraries (Supplementary Figure S7). Under these conditions, both new and archival poly(A) PCR preamplifications are compatible with RNA sequencing.

**Figure 6.**
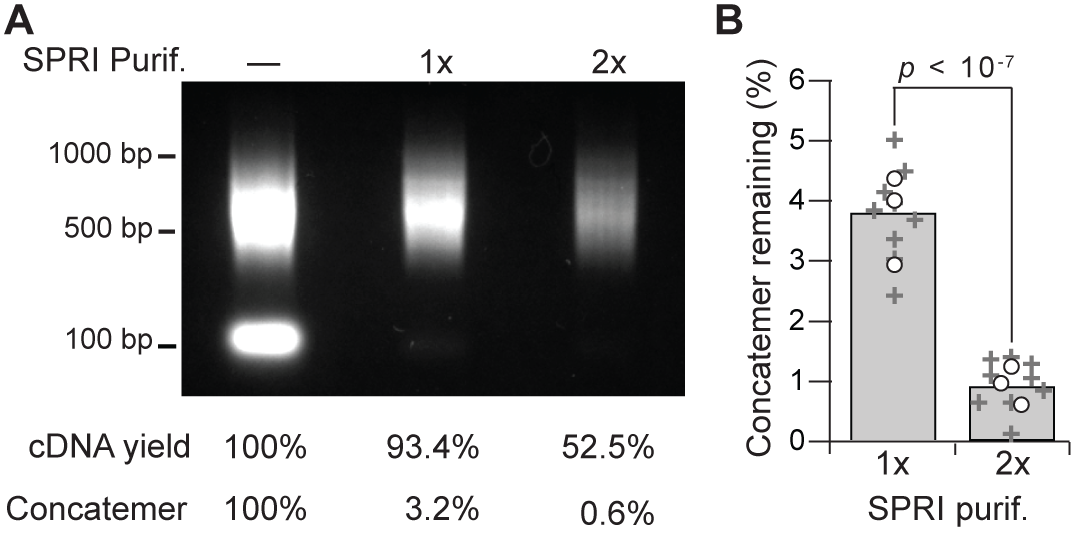
Iterative SPRI bead purification eliminates low molecular-weight contaminants before tagmentation. **(A)** Poly(A) PCR reamplifications (38) of 10-cell human breast cancer samples were analyzed by gel electrophoresis without purification or after one (1x) or two (2x) rounds of purification with 70% (vol/vol) SPRI beads. **(B)** Contaminating low molecular-weight concatemers are significantly reduced after two rounds of SPRI bead purification. Data are shown as the mean (gray) of *n* = 3 independent reamplifications (circles) each purified three times (+). Differences were assessed by two-way ANOVA with replication.

### Paired comparison of 10-cell transcriptomics by BeadChip microarray and RNA-seq

Poly(A) PCR provides an abundant source of material for transcript quantification, creating an opportunity to revisit 10-cell samples profiled earlier on BeadChip microarrays. In the original application of stochastic profiling, 10-cell samples were microdissected from 3D spheroids of a clonal human breast-epithelial cell line (38). We sequenced 18 biological replicates from this study (6.6 ± 2.3 million reads) along with three 10-cell pool-and-split controls that assessed technical variability (29,38). Technical correlation was as high within pool-and-split replicates measured by RNA-seq as when the same replicates were measured by microarray (*R* ~ 0.9; Figure 7B,C,D,F–H). For both platforms, undetectable genes in one technical replicate were quantified up to ~10^2^ = 100 transcripts per million (TPM) or ~10^3.3^ = 2000 BeadChip fluorescence intensity in another replicate. Among detected genes with at-least one technical replicate yielding zero measured TPM, we found that RNA-seq correlated with BeadChip intensity across replicates (*R* ~ 0.4, *p* ~ 0; Supplementary Figure S8A). The concordance between the two platforms strongly argues that transcript losses are authentic dropout events (73), not artifacts of RNA-seq read depth or BeadChip detection sensitivity. Combining the reliable detection limits of 100 TPM (Figure 7B,C,F) and ~250 ERCC copies/reaction (Figure 4B), we predict (250 copies/reaction)/(10 cells/reaction x 100 TPM) = 250,000 mRNA copies per cell, consistent with published estimates (35).

**Figure 7.**
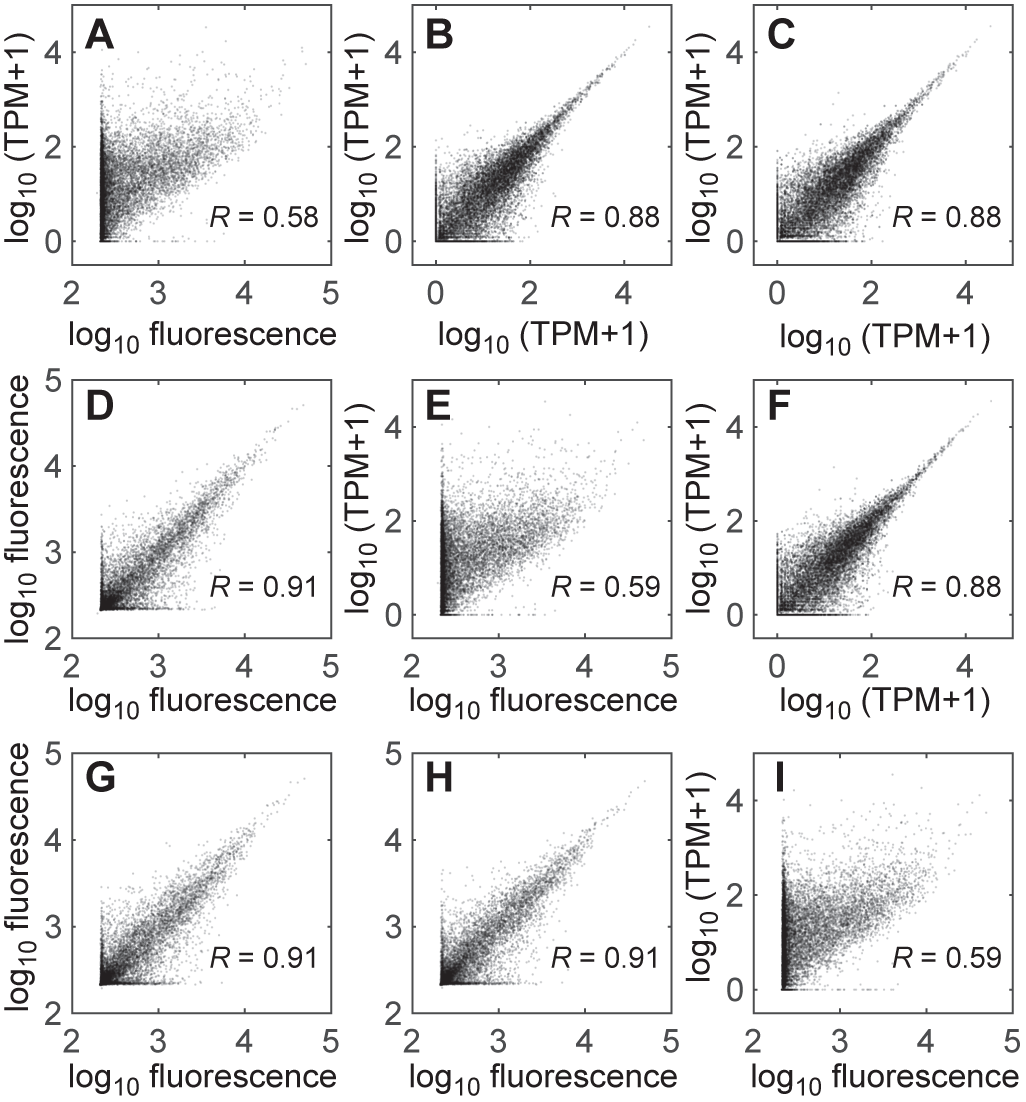
Paired comparison of 10-cell transcriptomes profiled by BeadChip microarray and 10cRNA-seq. **(A–I)** Three pool-and-split 10-cell replicates from before (38) were reamplified, purified, and tagmented for RNA-seq. Cross-correlations between replicates and measurement platforms are shown long with the log-scaled Pearson correlation (*R*).

When 10-cell transcript representation was compared, we found that RNA-seq TPM and BeadChip microarray intensities were correlated (*R* ~ 0.6; Figure 7A,E,I), albeit not as strongly as reported elsewhere (47,74). Some genes yielded background fluorescence on microarrays but moderate-to-high TPM, likely due to BeadChip probe sequences absent from the amplicons generated by poly(A) PCR. Among genes with a median TPM > 1000 by RNA-seq, we identified 27 BeadChip probes exhibiting a median fluorescence less than 10^2.5^. The median distance of the 27 probes from the 3’ end of the corresponding gene was 845 bases (IQR: 492–1392 bases), upstream of the distal ~500 bp 3’ ends amplified by poly(A) PCR. The probe-independent nature of RNA-seq reinforces one of its critical advantages for 10-cell transcriptomics.

We also evaluated quantitative concordance of the 18 10-cell samples measured both by BeadChip microarray and RNA-seq. The variance of 7713 genes was twice their mean value measured on each platform, suggesting significant biological variation across the 18 samples (*p* < 0.01). For biologically variable genes, the median sample-by-sample Pearson correlation between BeadChip microarray and RNA-seq was 0.42 (interquartile range: 0.16–0.63), with 599 transcripts showing R ≥ 0.8 (Supplementary Figure S8B). Considering a median TPM of 17 (interquartile range: 4–49) for the 10-cell data analyzed, these cross-platform correlations fall within the range reported for TCGA microarrays and RNA-seq (R ~ 0.4–0.9) (74). Our retrospective analysis indicates that 10cRNA-seq data corroborate BeadChip microarrays and provide broader access to 3’ mRNA ends not represented on oligonucleotide probe sets.

### Advantages of 10cRNA-seq for diverse mouse and human cell types

Last, we aggregated the intermediate revisions to 10-cell transcriptomic profiling (Figure 1) and asked whether there were more-overarching benefits to sequencing small pools versus single cells. Different methods for scRNA-seq have already been rigorously compared by multiple groups (45,69). Since a 10-cell approach could be adopted by many of these approaches, we focused instead on the data quality from published scRNA-seq datasets of various types relative to similar cells profiled by our 10cRNA-seq approach, including biological replicates and pool-and-split controls. We identified two scRNA-seq datasets for murine OPCs (75,76), two for murine lung neuroendocrine cells (77), two for human breast cancer (78,79), and one for MCF-10A cells (80). All raw data were identically processed and aligned to the transcriptome with RSEM (81). Using transcriptome references stringently emphasized exonic read alignments, and the RSEM model for expectation maximization enabled the degeneracy of 3’-end sequences to contribute to transcript quantification. Data quality was gauged by the percentage of reads aligned, and sensitivity was assessed by the number of Ensembl genes with an estimated TPM greater than one.

For the mouse cell types, we observed significant increases in gene detection between 10cRNA-seq and certain scRNA-seq datasets (Figure 8A). OPCs isolated by fluorescence-guided LCM showed better gene detection with 10cRNA-seq compared to scRNA-seq of OPCs purified by fluorescence-activated cell sorting (GSE75330) (76). Interestingly, gene detection in the sorted OPCs was also poorer than when OPCs were collected randomly in a cell atlas of the mouse cortex (GSE60361) (75), emphasizing the stresses caused by non-LCM methods of enrichment. We were unable to detect a significant increase in gene detection between small-cell lung cancer cells profiled by 10cRNA-seq and single neuroendocrine cells randomly dissociated from the mouse airway and profiled by plate-based scRNA-seq (77). However, neuroendocrine cells are so rare in this tissue that plate-based scRNA-seq was very underpowered (*n* = 5 cells). When droplet-based scRNA-seq was used to increase statistical power to *n* = 92 cells, there was a significant reduction in gene sensitivity compared to 10cRNA-seq profiling the equivalent of 120 cells (*n* = 12 10-cell replicates). In cases where gene sensitivities were comparable, we noted dramatically improved alignment rates for 10cRNA-seq (Figure 8B), reinforcing the efficiency of data collection by adopting a 10-cell approach.

**Figure 8.**
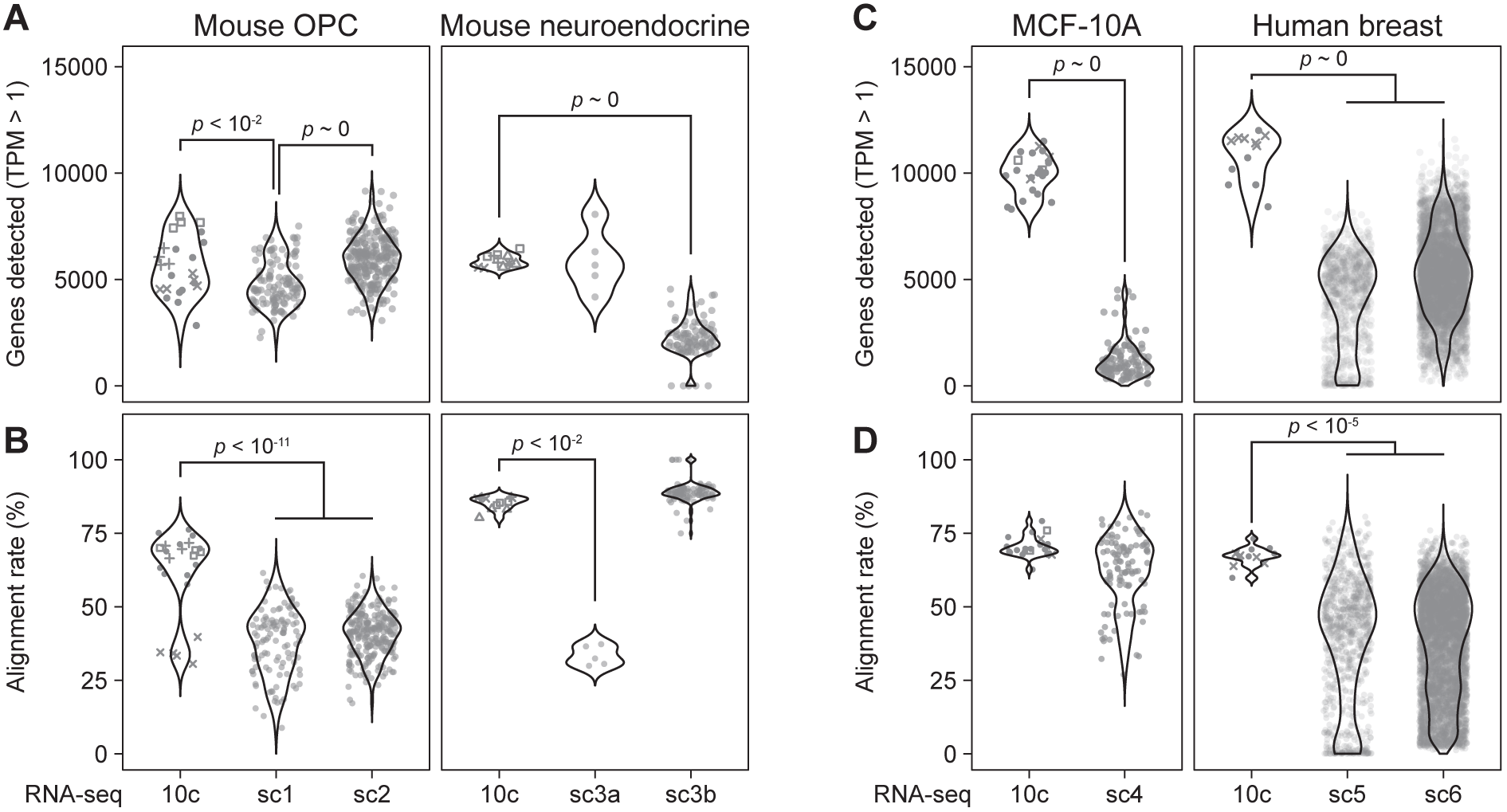
Improved gene detection and exonic alignment rates for 10cRNA-seq compared to scRNA-seq. **(A)** Detection of murine Ensembl genes for mouse oligodendrocyte precursor cells (OPCs) and lung neuroendocrine-derived cells. **(B)** Alignment rate comparison for OPCs and lung neuroendocrine-derived cells. **(C)** Detection of human Ensembl genes for MCF-10A cells and human breast cancer cells. **(D)** Alignment rate comparison for MCF-10A cells and human breast cancer cells. Public scRNA-seq data were obtained from the indicated accession numbers: sc1=GSE75330, sc2=GSE60361, sc3a=GSE103354 (plate-based), sc3b=GSE103354 (droplet-based), sc4=GSE66357, sc5=GSE113197, sc6=PRJNA396019. 10cRNA-seq data were aggregated from independent 10-cell samples (circles) and 10-cell equivalents from pool-and-split controls. Pool-and-split controls from the same day are indicated with non-circular markers corresponding to the shared day. Pairwise differences between 10-cell and single-cell methods were assessed by permutation test.

Results were similar but even more striking for human cell types (Figure 8C,D). 10cRNA-seq of MCF-10A cells and primary breast cancer cells routinely exceeded 10,000 Ensembl genes, the upper limit for any single cell profiled by three different scRNA-seq methods (78-80). Exonic alignment rates were also all significantly higher and comparable to the 10cRNA-seq alignments obtained with murine cells. The data suggest that mouse and human transcriptomes are sufficiently annotated to be used as reference alignments for limiting quantities of RNA, such as that extracted by LCM in situ for 10cRNA-seq.

## DISCUSSION

Single-cell transcriptomics has expanded or rewritten the catalog of cell types in tissues, organs, and organisms (77, 82-87). Yet, scRNA-seq does not obviate the need for complementary approaches, which accurately profile regulatory-state changes within a given cell lineage (40). The technical advances reported here demonstrate the immediate feasibility of 10cRNA-seq for mouse and human samples obtained in situ by LCM. We combined straightforward extensions of ERCC spike-ins and tagmentation with new approaches for fluorescence-guided LCM and cDNA purification that may prove beneficial for other applications (Figure 1). Although small-sample RNA-seq is never fully dissociated from tissue acquisition or cell handling, our data illustrate a workflow that can be paused and restarted when LCM is used as an intermediate step.

Previous descriptions of fluorescence-guided LCM relied upon fluorophores added by lectins, antibodies, or viruses (56,57,88). Through careful optimization of cryoembedding and LCM, we identified conditions that preserved the most-common fluorescent proteins used to engineer the mouse germ line. Compatibility with genomically encoded labels creates new opportunities for combining 10cRNA-seq with lineage tracing (89) to examine early regulatory-state changes in development and disease. Compared to fluorophore localization, RNA integrity was not as exquisitely sensitive to sample preparation and handling. Nevertheless, we recommend fresh cryoembedding of all samples in case other protein-guided approaches, such as immuno-LCM (90), might be pursued. The breast core biopsies profiled here were prospectively obtained and cryoembedded during an outpatient procedure. However, a nearly identical protocol has been deployed intra-operatively for surgical pathology (91), implying that fresh cryoembedding is not prohibitive for biobanked clinical samples.

A startling result from the revised protocol was the extent of poly(A) amplification observed in murine samples when reverse transcription was omitted. Nonspecific amplification was not as prominent in human samples obtained by LCM, pointing to specific differences in genome composition and the susceptibility to priming with AL1. A plausible explanation lies in transposable elements—specifically, the distinct classes of short interspersed nuclear elements (SINEs) in rodents and humans (92). Human-specific Alu SINEs and rodent-specific B-type SINEs both contain stretches of 10–20 As that could partially anneal to the T homopolymer sequence on the 3’ end of AL1 (93). However, to amplify during poly(A) PCR, an antisense SINE must be sufficiently nearby. The mouse genome is ~20% smaller than humans, and B-type SINEs are ~25% more numerous in mice compared to Alu SINEs in humans (92). The differences reduce the expected spacing of sense-antisense SINEs from ~6 kb in humans to ~4 kb in mice, consistent with a prior analysis of sense-antisense SINEs around transcription start sites (94). The shorter average spacing may be close enough for genomic fragments to compete with the ~500 bp cDNA amplicons generated during reverse transcription (Figure 3, 5A). Such nonspecific products were prevented from coamplifying with cDNA by using biotinylated oligo(dT)_24_ and streptavidin beads, akin to the bead capture and primer extension of droplet-based approaches (30,95). This strategy may prove useful in other non-murine settings, such as suspension cells, where genomic contamination will be more extensive than LCM (39).

ERCC spike-ins provide a standard to compare 10cRNA-seq against single-cell methods for transcriptomic profiling. Using the metrics of Svennson et al. (45), we estimate a 50% detection sensitivity of 45 copies per reaction (90% nonparametric CI: [15–485]) and a Pearson product-moment correlation coefficient of *R* = 0.86 (90% nonparametric CI: [0.71–0.91] from *n* = 72 samples). The *R* accuracy is somewhat lower than prevailing techniques, but that may be overly pessimistic because 10cRNA-seq uses such a dilute mix of spike-ins (4 million-fold dilution of the ERCC stock). Detection sensitivity is comparable to that reported for the most popular plate-based scRNA-seq methods, including SMART-seq2 (96) and CEL-seq (97). The strength of 10cRNA-seq lies in the use of 10-cell pooling to improve the per-cell sensitivity beyond the best microfluidic- and droplet-based approaches for scRNA-seq (45). Adopting a 10-cell approach may also prove beneficial for other approaches, such as the recent pairing of SMART-seq2 with LCM (34).

When 10cRNA-seq was compared to scRNA-seq, we often observed significant improvements in exonic alignment. Methods for scRNA-seq typically yield exonic alignment rates below 50% (80), with the remainder of aligned reads splitting equally between intronic and intergenic sequences (96). 10cRNA-seq achieves exonic alignments of 70% or higher despite using oligo(dT)-primed reverse transcription with the same potential to prime internal A homopolymer sequence as with scRNA-seq (98,99). Interestingly, in one instance of similarly high exonic alignment (GSE66357, Figure 8B), the RNA-printing approach to scRNA-seq incorporated a DNase treatment absent from all other methods (80). This study also yielded a significantly reduced gene-detection sensitivity compared to 10cRNA-seq. Commingling genomic DNA may dilute exonic alignment percentages and inflate the number of genes detected due to chance sequencing of genomic DNA from exonic loci. Multiple scRNA-seq approaches incorporate unique molecular identifiers appended to oligo(dT) (45,80,100). The identifiers avoid redundantly counting the same product of reverse transcription, and they also retrospectively exclude sequenced reads that do not come from cDNA. The biotin cleanup approach we devised for mouse cells (Figure 5) achieves cDNA selection prospectively in situations where genomic contamination may be problematic.

Our work illustrates that 10-cell profiling can extend beyond microarrays (42) and quantitative PCR (36,37) to compete favorably with scRNA-seq. Although ill-suited for lineage mapping of highly mixed cell populations (40), 10cRNA-seq exploits the precision of LCM to target specific cell types in situ and define their regulatory heterogeneities. LCM is also advantageous for sequencing cells that are delicate or difficult to dissociate rapidly (34). We anticipate immediate applications of 10cRNA-seq to cancer biology, where the initiation, progression, and diversification of tumors could be tackled in modern animal models as well as in patients.

## ACCESSION NUMBERS

All 10cRNA-seq data are available through the NCBI Gene Expression Omnibus (GSE120261).

## SUPPLEMENTARY DATA

Supplementary Data are available at NAR online.

## ACKNOWLEDGEMENT

We thank Kathy Repich and Jennifer Harvey for help with clinical sample acquisition, Emily Farber for technical assistance with RNA sequencing, and Stephen Turner from the UVA Bioinformatics Core for guidance on alignment approaches.

